# Immune-induced remodelling of mRNA structurome regulates uORF-mediated translation

**DOI:** 10.1101/2023.05.09.540021

**Authors:** Yezi Xiang, Tianyuan Chen, Patrick S. Irving, Kevin M. Weeks, Xinnian Dong

## Abstract

To survive stress, eukaryotes selectively translate stress-related transcripts while inhibiting growth-associated protein production. How this translational reprogramming occurs under biotic stress has not been systematically studied. To identify common features shared by transcripts with stress-upregulated translation efficiency (TE-up), we first performed high-resolution ribosome-sequencing in *Arabidopsis* during pattern-triggered immunity and found that TE-up transcripts are enriched with upstream open reading frames (uORFs). Under non-stress conditions, start codons of these uORFs (uAUGs) have higher-than-background ribosomal association. Upon immune induction, there is an overall downshift in ribosome occupancy at uAUGs, accompanied by enhanced translation of main ORFs (mORFs). Using *in planta* nucleotide-resolution mRNA structurome probing, we discovered that this stress-induced switch in translation is mediated by highly structured regions detected downstream of uAUGs in TE-up transcripts. Without stress, these structures are responsible for uORF-mediated inhibition of mORF translation by slowing progression of the translation preinitiation complex to initiate translation from uAUGs, instead of mAUGs. In response to immune induction, uORF-inhibition is alleviated by three Ded1p/DDX3X-homologous RNA helicases which unwind the RNA structures, allowing ribosomes to bypass the inhibitory uORFs and upregulate defence protein production. Conservation of the RNA helicases suggests that mRNA structurome remodelling is a general mechanism for stress-induced translation across kingdoms.

## MAIN

In eukaryotes, protein translation is normally cap-dependent. The m^7^G-cap of the mRNA is recognized by the 43S translation preinitiation complex comprised of the 40S ribosomal subunit and the eIF2-GTP-Met-tRNAi ternary complex. The preinitiation complex then scans along the 5’ leader sequence of the mRNA to initiate translation at a start codon^1–3^. It has been proposed that ribosome scanning and start codon selection are regulated by elements in the 5’ leader sequence, such as RNA primary sequences (for example, the Kozak sequence context), upstream open reading frames (uORFs), secondary structures, and RNA modifications^4–7^. Under stress conditions, such as nutrition depletion^8^, hypoxia^9, 10^, or pathogen challenge^11^, global translation is reprogrammed, leading to elevated stress-responsive protein production, but repressed growth-related protein synthesis, which is crucial to the survival and adaptation to stress. The mechanisms involved in the stress-induced translation have been investigated for a small number of key transcription factors (for example, yeast general control nondepressible 4 (GCN4)^12^ and mammalian activating transcription factor 4 (ATF4)^13^), whose translation is normally inhibited by the uORFs in the 5’ leader sequences of their mRNAs. Upon stress, phosphorylation of eukaryotic Initiation Factor 2α (eIF2α) decreases the available ternary complex, resulting in reduced translation initiation from the start codons of uORFs (uAUGs) and prolonged scanning of the preinitiation complex to translate the downstream main open reading frames (mORFs) to promote cell survival^12–15^. However, this integrated stress response pathway has been shown to be dispensable in plant responses to biotic and abiotic stresses studied so far^11, 16–18^. Moreover, recent global ribosome-sequencing (Ribo-seq, sequencing of ribosome-protected RNA fragments) studies have shown that uORFs are a prevalent feature in eukaryotic mRNAs, not limited to these few well-studied examples^19–21^. This raises questions: How is translation of these uORF-containing mRNAs regulated if not by phosphorylation of eIF2α? Are there common features in transcripts that are responsible for stress-induced translation?

### Global ribosome dynamics on uORFs and mORFs during pattern-triggered immunity

To identify novel regulatory mechanisms involved in uORF-mediated translation, we performed high resolution Ribo-seq on *Arabidopsis* seedlings in response to the induction of pattern-triggered immunity by elf18 (N-terminal epitope of the bacterial elongation factor Tu)^22^ to enable us to examine the translational activities in 5’ leader sequences (see Extended Data Fig. 1 and Methods for details). Comparing the elf18-treated samples to mock-treated controls, we identified, among the 13051 expressed transcripts, 1157 with increased translational efficiency (TE-up) and 1150 with decreased translational efficiency (TE-down) (Fig. 1a and Extended Data Fig. 2a, b). Gene ontology (GO) analysis^23–25^ of the TE-up genes revealed an enrichment of biological processes in response to a variety of environmental stresses, such as biotic stimulus, abiotic stimulus, and chemicals, whereas GO terms for the TE-down genes are mostly growth-related metabolic processes (Extended Data Fig. 2c, d).

Next, we asked whether there are intrinsic sequence features in TE-up mRNAs that attribute to their selective translation during the immune response by searching for uAUGs with translation activities based on their above-background levels of ribosome occupancy (see Extended Data Fig. 2a, b and Methods for details). We found that compared to the percentage in all expressed transcripts (21.8% of the transcripts), “translating uAUGs”, which can appear one or more times per transcript, are significantly enriched in TE-up transcripts (30%), but depleted in TE-down mRNAs (16.7%) (Fig. 1b). This finding suggests that translation initiation from uAUGs may impact translation of downstream mORFs during the immune response.

**Fig. 1.**
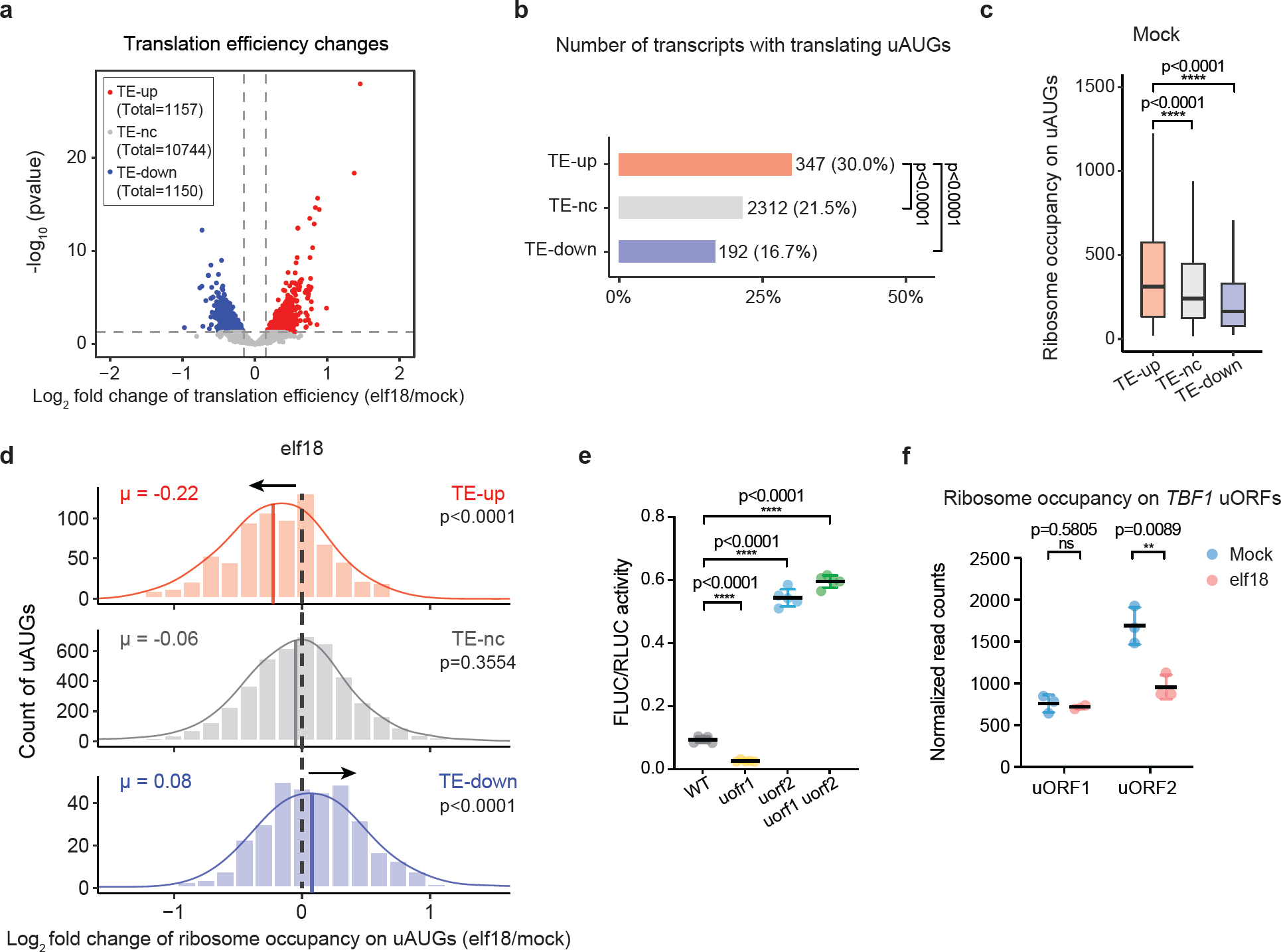
Translational dynamics of uORF-containing transcripts during pattern-triggered immunity. **a**, A volcano plot of global translational efficiency (TE) changes during pattern-triggered immunity. TE-up: transcripts with upregulated TE (*p* value < 0.05, log_2_ fold change > 0.16); TE-nc: transcripts with no changes in TE (*p* value > 0.05); TE-down: transcripts with downregulated TE (*p* value < 0.05, log_2_ fold change < -0.16). **b**, Number and percent of transcripts with translating uAUGs in the TE-up, TE-nc, and TE-down groups. Fisher’s Exact Test was used to determine the *p* value of the difference between groups. **c**, A box plot of ribosome occupancy (normalized read counts) on translating uAUGs in the TE-up, TE-nc, and TE-down transcripts under the mock condition. Boxes represent the interquartile range (IQR), and whiskers indicate data within 1.5 × IQR of the top (Q3) and bottom (Q1) quartiles. *P* values were calculated by two tailed Mann-Whitney tests. **d**, Histograms with density curves of log_2_ fold change of ribosome occupancy on translating uAUGs in the TE-up, TE-nc, and TE-down transcripts in response to elf18 treatment. μ, averaged log_2_ fold change value. *P* values were calculated by two tailed paired t tests. **e**, Mutagenesis of start codons of uORF1 and/or uORF2 in the *TBF1* 5’ leader sequence and translational activity in the *35S:TBF1 5’ leader sequence-FLUC/35S:RLUC* dual-luciferase reporter assay. Data were analysed by two tailed student’s t test. Values are means ± SDs. **f**, Ribosome occupancy on uORF1 and uORF2 in the endogenous *TBF1* transcript in response to elf18 treatment.

We then specifically examined the ribosome dynamics on the 763 translating uAUGs in the TE-up transcripts and found that under mock condition, they have significantly higher ribosome association compared to the 5626 translating uAUGs in all expressed transcripts (Fig. 1c). In response to elf18 treatment, we detected an overall decrease in ribosome occupancy on the uAUGs of TE-up transcripts (Fig. 1d) and this reduction is even more striking for the subset of transcripts with only one translating uAUG. As translation initiation from uAUGs typically inhibits the translation of downstream mORFs^6, 26^, this elf18-triggered reduction in ribosome association with uAUGs suggests an immune-induced release of uORF-mediated inhibition on downstream mORF translation.

### uORF-mediated regulation of the *TBF1* mRNA translation

To understand how translation initiation from uAUGs modulates the downstream protein production, we examined the translation regulation of an immune-responsive mRNA encoding the transcription factor TL1-binding factor (TBF1)^18^. The rapid and transient induction of *TBF1* mRNA translation during pattern-triggered immunity is controlled, in part, by two translating uORFs (namely uORF1 and uORF2)^18, 27^. To examine how uORF1 and uORF2 regulate downstream TBF1 translation, we used a dual-luciferase system in which translation of the constitutively transcribed firefly luciferase reporter (FLUC) is driven by the 5’ leader sequence of *TBF1* and the resulting reporter activity is normalized to that of the constitutively expressed Renilla luciferase (RLUC) on the same construct. We then performed mutagenesis on the *TBF1* 5’ leader sequence to determine how changes in uORFs influence the translation of downstream FLUC by transiently expressing wild type (WT) or the mutant dual-luciferase reporters in *Nicotiana benthamiana* (*N. benthamiana*). We found that mutating the start codon of uORF1 (uAUG1) to CUG reduced the FLUC activity, whereas mutating the start codon of uORF2 (uAUG2) or both uAUG1 and uAUG2 to CUG caused a significant increase in FLUC translation (Fig. 1e). These findings indicate that protein production from mORF is positively regulated by uORF1 translation, but significantly inhibited by uORF2 translation.

We next examined the ribosome dynamics on uORF1 and uORF2 during pattern-triggered immunity using the Ribo-seq data. We observed a significant reduction in ribosome occupancy on uORF2 in response to elf18 treatment, but no change for uORF1 (Fig. 1f). Given the inhibitory role of uORF2 in regulating the translation of TBF1, our data suggest that elf18-induced downshift in ribosome occupancy on the inhibitory uORF2 reflects the release of uORF2 inhibition on mORF translation, consistent with what we observed on the global scale for all elf18-induced TE-up transcripts (Fig. 1d).

### *In planta* SHAPE-MaP of RNA secondary structurome changes during pattern-triggered immunity

To examine the mechanisms that underly the immune-induced switch in ribosome occupancy from uORFs to mORFs in the TE-up transcripts, we first assessed the Kozak sequence context flanking the AUGs, because this -3 to +4 (with A in AUG being +1) region could impact the start codon recognition by the translation preinitiation complex^5^. Even though a Kozak sequence context has not been defined through a comprehensive study of plant mRNAs, we analysed the AG-content of the Kozak contexts of uAUGs and mAUGs in all the expressed transcripts because A and G have been shown to support higher translational activity in a previous study^28^, an observation that is confirmed by our own Ribo-seq data (Extended Data Fig. 3). We found that mAUGs have remarkably higher AG-contents than translating uAUGs (Fig. 2a), in agreement with recent findings in animals that uAUGs tend to have less preferable Kozak sequence contexts compared to mAUGs^29, 30^. We next compared the Kozak contexts for translating uAUGs among the TE-up, TE-nc, and TE-down transcripts, and found that they have similar AG-contents (Fig. 2a). This suggests that while the Kozak sequence context is important for start codon recognition under static conditions, it is unlikely to be responsible for the elf18-mediated switch from uAUG to mAUG translation in the TE-up transcripts.

**Fig. 2.**
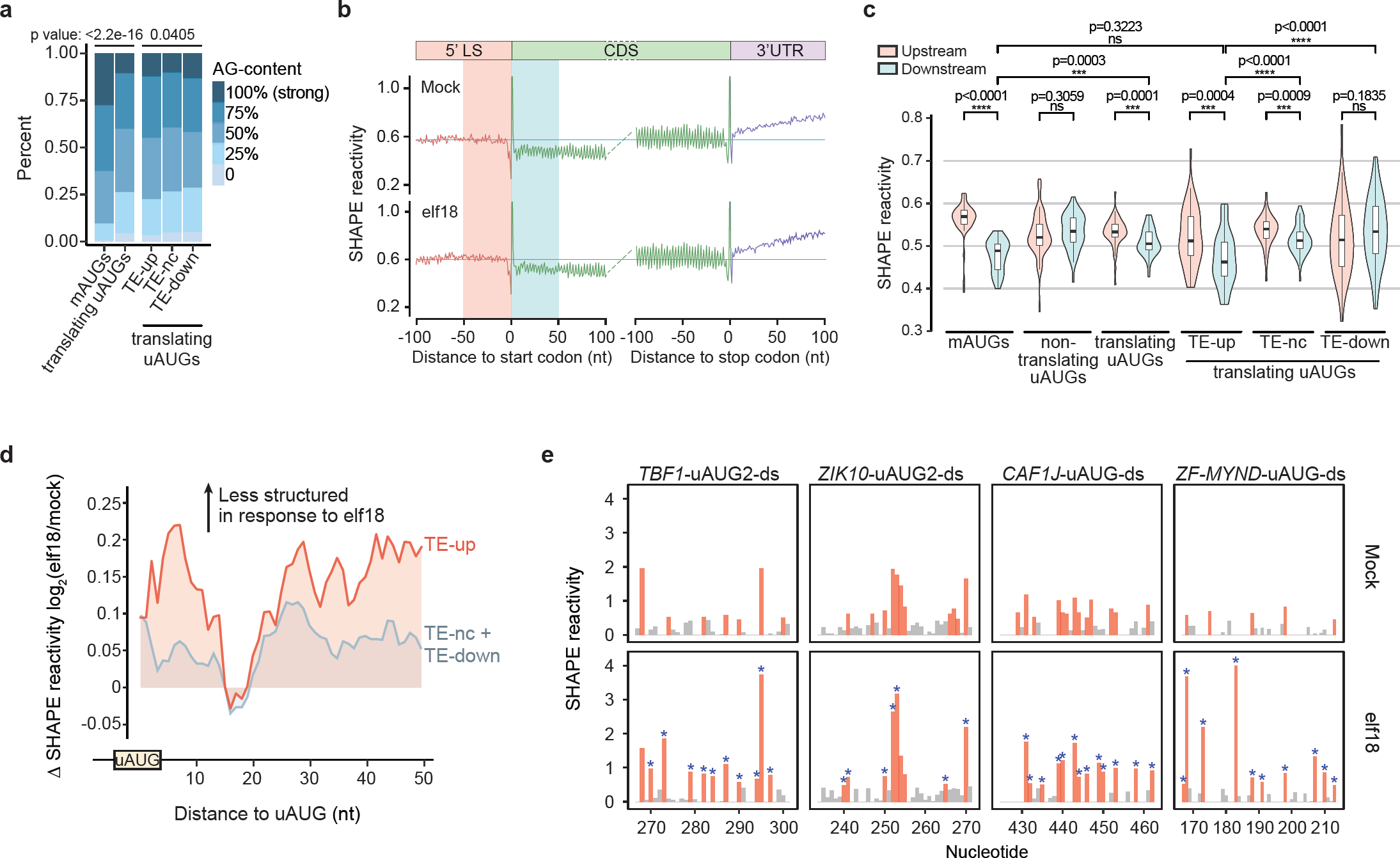
Global SHAPE-MaP of *in planta* RNA secondary structural remodelling during pattern-triggered immunity. **a**, Comparison of the Kozak sequence contexts (AG-content) flanking mAUGs and translating uAUGs. *P* values were calculated by Chi-Square test. **b**, Averaged SHAPE reactivities in the 5’ leader sequence, CDS, and 3’ UTR across all transcripts aligned by the start and stop codons of mORFs. **c**, Violin plots show the comparisons of SHAPE reactivities of the 50 nt upstream and the 50 nt downstream of mAUGs and uAUGs in different categories of transcripts under the mock condition. **d**, elf18-induced averaged SHAPE reactivity changes across nucleotide downstream of translating uAUGs in TE-up transcripts (red) or TE-nc and TE-down transcripts (grey). **e**, SHAPE reactivity changes in four representative TE-up transcripts that contain translating uAUGs. uAUGs are highlighted by arrows and SHAPE reactivities of the downstream structured regions are highlighted in light blue. Orange and red bars are nucleotides with median to high SHAPE reactivities. Blue asterisks, nucleotides with elf18-induced increases in SHAPE reactivities.

We next investigated immune-induced mRNA secondary structural changes by adapting <u>s</u>elective 2’-<u>h</u>ydroxyl <u>a</u>cylation and <u>p</u>rimer <u>e</u>xtension based on the <u>m</u>utational <u>p</u>rofile (SHAPE-MaP), which has been successfully applied in *Escherichia coli*^31^ and mammalian cells^32–34^, to detect *in planta* RNA secondary structurome changes at nucleotide resolution. This strategy relies on SHAPE reagents^35, 36^ (here, 2-methylnicotinic acid imidazolide, NAI), a group of hydroxyl-selective electrophiles that react with the 2’-hydroxyl position of unpaired residues of RNA regardless whether they are associated with RNA-binding proteins^37, 38^ (Extended Data Fig. 4a). The resulting 2’-O-adducts cause mutations in the cDNA during reverse transcription. DNA libraries are then sequenced and sites of mutations are aligned to RNA sequences to create SHAPE reactivity profiles, which provide a per-nucleotide quantification of RNA structures. In this analysis, higher SHAPE reactivity indicates less RNA structural complexity. To validate our protocol, we first performed targeted *in planta* SHAPE-MaP (see Methods for details) to probe the secondary structure of *Arabidopsis* 18S rRNA, which had been studied by Ding et al.^39^ using a DMS-based structure-seq method. Our results were not only consistent with those found by the structure-seq (Extended Data Fig. 4b, blue), but also uncovered additional unpaired regions with higher percentage of G and U (Extended Data Fig. 4b, orange), confirming that SHAPE-MaP measures flexibility at all four ribonucleotides, without the detection bias towards A and C of the conventional DMS-based probing^33^.

We then performed the global *in planta* SHAPE-MaP analysis of mRNAs in *Arabidopsis* seedlings without and with elf18 treatment. The quality of the data was confirmed by a strong correlation among the data obtained from all four replicates under each of the conditions (Extended Data Fig. 4c) and an overall higher mutation rate in all four nucleotides in the NAI-modified samples than that in the unmodified control samples (Extended Data Fig. 4d). For subsequent analyses, only data that passed the stringent cut-offs for read depth and completeness were used to enable accurate structure modelling (see Methods for details)^40^. We found that in both mock- and elf18-treated samples, the cumulative SHAPE reactivities of the nucleotides in the 5’ leader sequences and coding sequences (CDSs) across the 13051 expressed transcripts were modestly lower than those in the 3’ untranslated regions (3’ UTRs) (Extended Data Fig. 4e). As observed in the structuromic analyses conducted in other studies^39, 41^, there was an extraordinarily high average SHAPE reactivity at the mAUGs and the stop codons of CDS, and a strong three-nucleotide structure periodicity across CDSs (Fig. 2b).

Interestingly, we noticed that although the overall SHAPE reactivities of the 5’ leader sequences and the CDSs are comparable, the region immediately downstream of the mAUGs had a lower average SHAPE reactivity (Fig. 2b, c), indicating a higher structural complexity in the region. We wondered whether this structural feature is related to start codon selection and translation initiation from mAUGs as proposed in a previous reporter study using an artificial hairpin structure^42^; and whether a similar feature exists for uAUGs. To answer these questions, we examined the SHAPE reactivities for each nucleotide in the regions 50 nucleotides (nt) upstream and downstream of AUGs. We found that nucleotides downstream of mAUGs and translating uAUGs had significantly lower SHAPE reactivities compared to those upstream, but this was not observed for non-translating uAUGs (Fig. 2c). We further investigated whether the observed feature may also contribute to the dynamic regulation of uAUG-mediated translation in the TE-up, TE-nc, and TE-down transcripts (Fig. 1c, d). We found that, under the mock condition, translating uAUGs in the TE-up transcripts had significantly lower SHAPE reactivities in their downstream regions compared to those in the TE-nc and TE-down transcripts (Fig. 2c). This may explain the inhibitory effect of uORFs on mORF translation in the TE-up transcripts in the absence of an immune signal, because RNA secondary structure downstream of uAUGs (here we name “uAUG-SS”) may slow the scanning of the translation preinitiation complex to enhance ribosome assembly (Fig. 1c) and initiate translation from uAUGs. As our Ribo-seq data detected an elf18-triggered decrease in ribosomal occupancy at the translating uAUGs of the TE-up transcripts (Fig. 1d), we hypothesize that elf18-induced translation in these transcripts may result from changes in uAUG-SS. Indeed, we observed a significant elf18-induced increase in average SHAPE reactivity in uAUG-SS of the TE-up transcripts (Fig. 2d), from which four examples with high read depth (among top 5%) are shown (Fig. 2e). We propose that the reduction in mRNA secondary structure downstream uAUGs allows the preinitiation complex to scan beyond the uAUGs to initiate translation from downstream mAUGs. Interestingly, a similar increase in average SHAPE reactivity was not observed for mAUG-SS of the TE-up transcripts (Extended Data Fig. 4g).

### RNA secondary structural changes regulate *TBF1* translation

To further explore how RNA secondary structures regulate translation, we selected the *TBF1* transcript for detailed examination. First, we performed a targeted *in planta* SHAPE-MaP on the endogenous *TBF1* 5’ leader sequence and found that under non-stress conditions, the region 35 nt downstream of uAUG2 is highly structured (Fig. 3a, b), consistent with what we observed globally for uAUGs in the TE-up transcripts (Fig. 2c). This highly structured region was also detected in the mRNA of the *TBF1* 5’ leader sequence-driven dual-luciferase reporter that was transiently overexpressed in *N. benthamiana* (Extended Data Fig. 5a), suggesting a similar folding pattern in *Arabidopsis* as well as in tobacco. In response to elf18 treatment, there was a significant increase in SHAPE reactivity and opening of the structure in this downstream region of uAUG2 (Fig. 3a, c and d), correlated with the decrease in ribosome association observed in our Rib-seq (Fig. 1f).

**Fig. 3.**
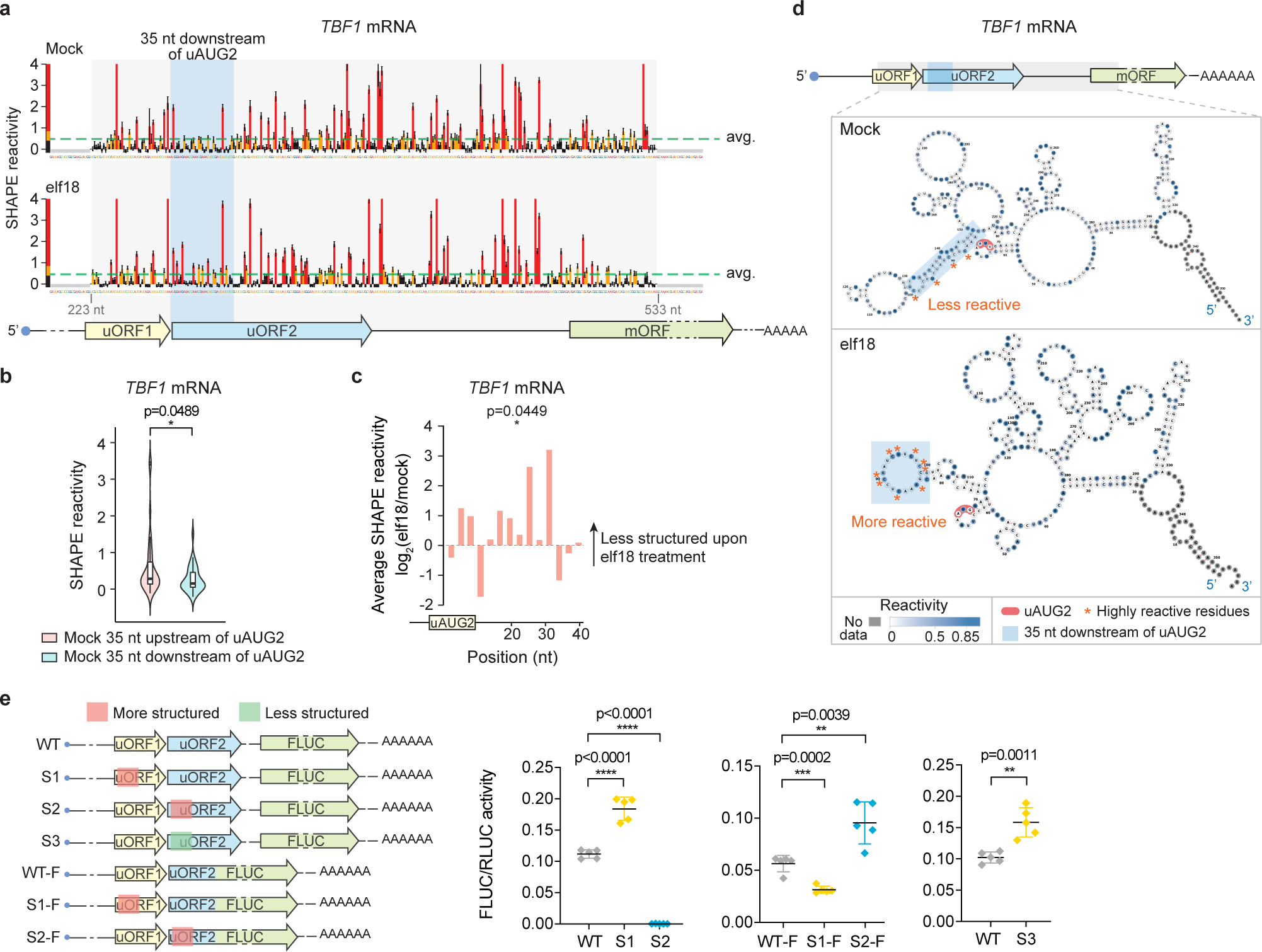
Immune-induced RNA secondary structure remodelling in the *TBF1* 5’ leader sequence regulates mORF translation. **a**, Overview of the SHAPE reactivities around uORF2 and mAUG of the endogenous *TBF1* transcript in *Arabidopsis* under mock (top) and elf18 treatment (bottom). The region 35 nt downstream of uAUG2 was highlighted in light blue. Green dashed lines indicate average SHAPE reactivity across the region. **b**, Comparison of SHAPE reactivities in the region 35 nt upstream and downstream of uAUG2 in the *TBF1* transcript under mock condition. Data were analysed by two tailed Mann-Whitney tests. **c**, elf18-induced average SHAPE reactivity changes calculated for every three nucleotides downstream of uAUG2. P-value shows the difference of SHAPE reactivities in the region 35 nt downstream of uAUG2 under mock and elf18 treatment (analysed by Wilcoxon matched-pairs signed rank tests). **d**, Modelling of the RNA secondary structures around uORF2 and mAUG under mock and elf18 treatment. The region 35 nt downstream of uAUG2 is highlighted in light blue. Residues with high reactivities are highlighted by orange asterisks. **e,** Effects of mutagenesis in the regions downstream of uAUG1 and uAUG2. WT-F, S1-F, and S2-F are uORF2 in-frame fusion with FLUC. Data were analysed by two tailed student’s t test. Values are means ± SDs.

We next performed mutagenesis on the *TBF1* 5’ leader sequence in the dual-luciferase system (Fig. 3e, WT) to determine if alterations of *TBF1* 5’ leader sequence secondary structures could influence the translation of downstream FLUC. We found that when we made the region downstream of uAUG1 more structured (Extended Data Fig. 5b, c), there was a significant increase in FLUC activity (Fig. 3e, S1). But when we made the region downstream of uAUG2 more structured, the FLUC activity was diminished (Fig. 3e, S2). These data are consistent with our observation that uORF1 positively regulates, while uORF2 significantly inhibits, the downstream FLUC translation (Fig. 1e) and support our hypothesis that a complex structure downstream of uAUGs facilitates their translation initiation by slowing the scanning of the translation preinitiation complex. To further demonstrate that the observed mutant effects on the mORF-FLUC reporter was caused by uORF2 translation, not by amino acid changes in the uORF2 peptide, nor structure-induced ribosome dissociation, we introduced the same structural mutations into a new reporter that has FLUC fused in-frame with only the first 45 nt of uORF2 (Fig. 3e, WT-F) to directly detect the translation of uORF2. As expected, when the region downstream of uAUG1 was mutated to be more structured, there was a decrease in the uORF2 reporter translation (Fig. 3e, S1-F) whereas when the region downstream of uAUG2 was mutated to be more structured, the reporter translation was increased (Fig. 3e, S1-F). Lastly, to mimic the structural changes in response to elf18, we mutated the 15-30 nt region downstream of uAUG2 to be less structured in the mORF-FLUC reporter (Extended Data Fig. 5b, c, S3) and observed a significant increase in the FLUC activity (Fig. 3e, S3). Altogether, these results demonstrate that a complex structure downstream of uAUGs (uAUG2 for the *TBF1* transcript) is conducive to uORF-mediated inhibition of downstream mORF translation by facilitating translation initiation from the uAUGs. This inhibition may be alleviated during stress when the RNA secondary structure is unwound to facilitate the translation preinitiation complex scanning beyond uAUGs to initiate mORF translation.

### Inducible RNA helicases unwind complex RNA structures and facilitate mORF translation

We next searched for RNA helicases that may play a role in unwinding uAUG-SS to facilitate stress-induced translation. To identify potential candidates, we examined the translational efficiency changes of the 54 known RNA helicases in *Arabidopsis* and found four candidates showing significant translational induction in response to elf18 (Fig. 4a). Among them, only RH37 is predicted to be localized in the cytoplasm. A genome-wide homology analysis across angiosperms revealed another two close RH37 homologues, RH11 and RH52, consistent with the result of a recent gene family study^43^. Through comparisons of protein sequences, functional domains and structures predicted by AlphaFold^44^ (Extended Data Fig. 6), we showed that these three helicases are potential homologues of the yeast Ded1p and the human DDX3X. The sequence and structural homology to the yeast Ded1p also aligns well with the anticipated function for RH11, RH37 and RH52, because mutating the yeast Ded1p helicase activity has been shown to cause an increase in translation initiation from near-cognate start codons immediately upstream of a structured region, resulting in inhibition of mORF translation^45^. Conversely, elf18-induced increases in RH11, RH37 and RH52 levels or activities could promote the unwinding of uAUG-SS, thus alleviating the uORF-inhibition on mORF translation.

**Fig. 4.**
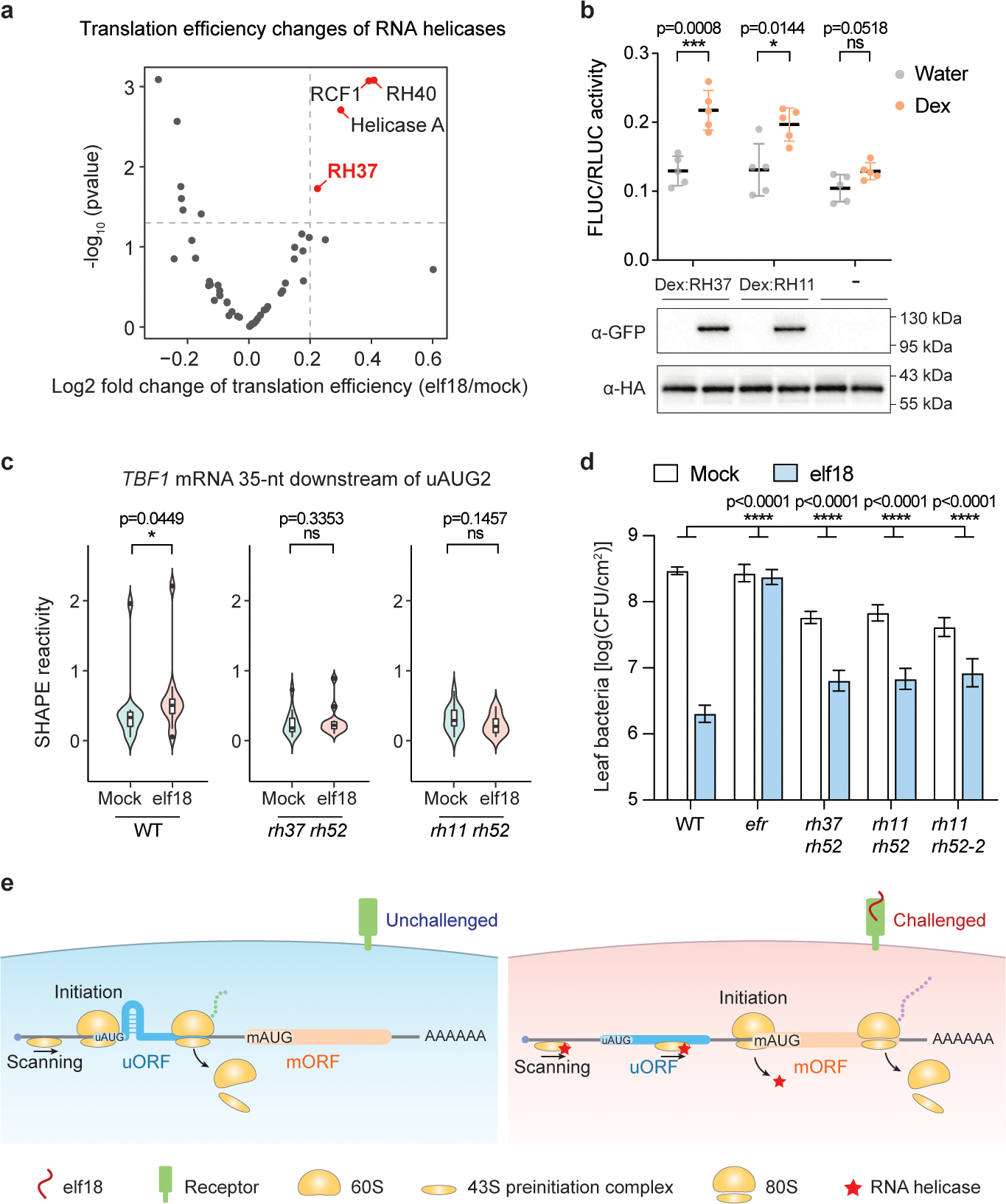
RNA helicases unwind RNA secondary structures to induce pattern-triggered immunity. **a**, A volcano plot of translational efficiency changes of 54 known RNA helicases upon elf18 treatment. **b**, Impact of Dex-induced expression of YFP-tagged RNA helicases (RH37, RH11) (bottom) in regulating translation of the *35S:TBF1 5’ leader sequence-FLUC/35S:RLUC* dual-luciferase reporter (top). HA-tagged RLUC levels were detected as internal controls. Data were analysed by two tailed student’s t test. Values are means ± SDs. **c**, Violin plots of *in planta* SHAPE reactivity changes in the region 35 nt downstream of *uAUG2* of the endogenous *TBF1* mRNA in wild type (WT) and helicase mutants. Boxes represent the interquartile range (IQR), and whiskers indicate data within 1.5 × IQR of the top (Q3) and bottom (Q1) quartiles. Circles represent values for outliers. *rh37 rh52* and *rh11 rh52*, helicase mutants. Data were analysed by Wilcoxon matched-pairs signed rank tests. **d**, The elf18-induced protection against *Psm* ES4326 on WT and helicase mutants (n ≥ 12 biological replications). Bacterial growth was measured at 2 days post inoculation and represented by means ± s.e.m. Data were analysed by two-way ANOVA. The experiment was repeated twice. **e**, A model on RNA secondary structure-mediated translational regulation of uORF-containing transcripts during pattern-triggered immunity.

To test our hypothesis, we built the constructs *Dex:RH37-YFP* and *Dex:RH11-YFP* to put the transcription of *RH37-YFP* and *RH11-YFP* under the control of a dexamethasone (dex)-inducible system^46^ and transiently coexpressed them in *N. benthamiana* with the dual-luciferase reporter driven by the 5’ leader sequence of *TBF1*. Strikingly, we observed a significant increase in the FLUC activities four hours after treatment with dex (Fig. 4b). This suggests that a transient increase in the expression of these RNA helicases could lead to enhanced translation of TBF1.

To demonstrate that RH11, RH37, and RH52 helicases are responsible for the elf18-induced unwinding of the region downstream of inhibitory uAUGs, we generated *rh37 rh52*, *rh11 rh52*, and *rh11 rh37* double mutant lines using a high efficiency CRISPR method^47^. Since the *rh11 rh37* mutant exhibited a developmental defect, whereas the *rh37 rh52* and the *rh11 rh52* plants displayed near WT morphology, we chose to use *rh37 rh52* and *rh11 rh52* for targeted *in planta* SHAPE-MaP of the endogenous *TBF1* transcript. We found that elf18-induced structural changes in the region 35 nt downstream of uAUG2 observed in WT are diminished in the double mutants (Fig. 4c), supporting the involvement of RH11, RH37, and RH52 in elf18-induced unwinding of uAUG-SS. It is worth noting that this group of helicases has been found to be associated with the translation preinitiation complex, instead of the 80S ribosome^45, 48^, consistent with our observation that the elf18-induced SHAPE reactivity increase in the downstream regions of uAUGs was not detected in the downstream regions of mAUGs (Extended Data Fig. 4g).

To examine the global effect of the helicase mutations on elf18-induced resistance, we performed bacterial infection using *Pseudomonas syringue* pv*. maculicola* ES4326 (*Psm* ES4326) in WT, *rh37 rh52* and *rh11 rh52* mutants after pre-treating plants with elf18. As a negative control, we included the elf18 receptor mutant, *efr*. We found that the helicase mutants have higher basal resistance to *Psm* ES4326 than the WT plants, suggesting that they affect transcripts other than those involved in pattern-triggered immunity (Fig. 4d). Regardless of this caveat, the specific defect of the mutants in translating mRNAs involved in pattern-triggered immunity is clearly demonstrated by their diminished sensitivity to elf18-induced resistance against *Psm* ES4326 infection (Fig. 4d). Altogether, our results demonstrate that the elf18-inducible RNA helicase RH37 and its homologues RH11 and RH52 are involved in unwinding of uAUG-SS in TE-up transcripts and translation of downstream defence proteins against pathogen challenge.

## Extended Data Figures

**Extended Data Fig. 1.**
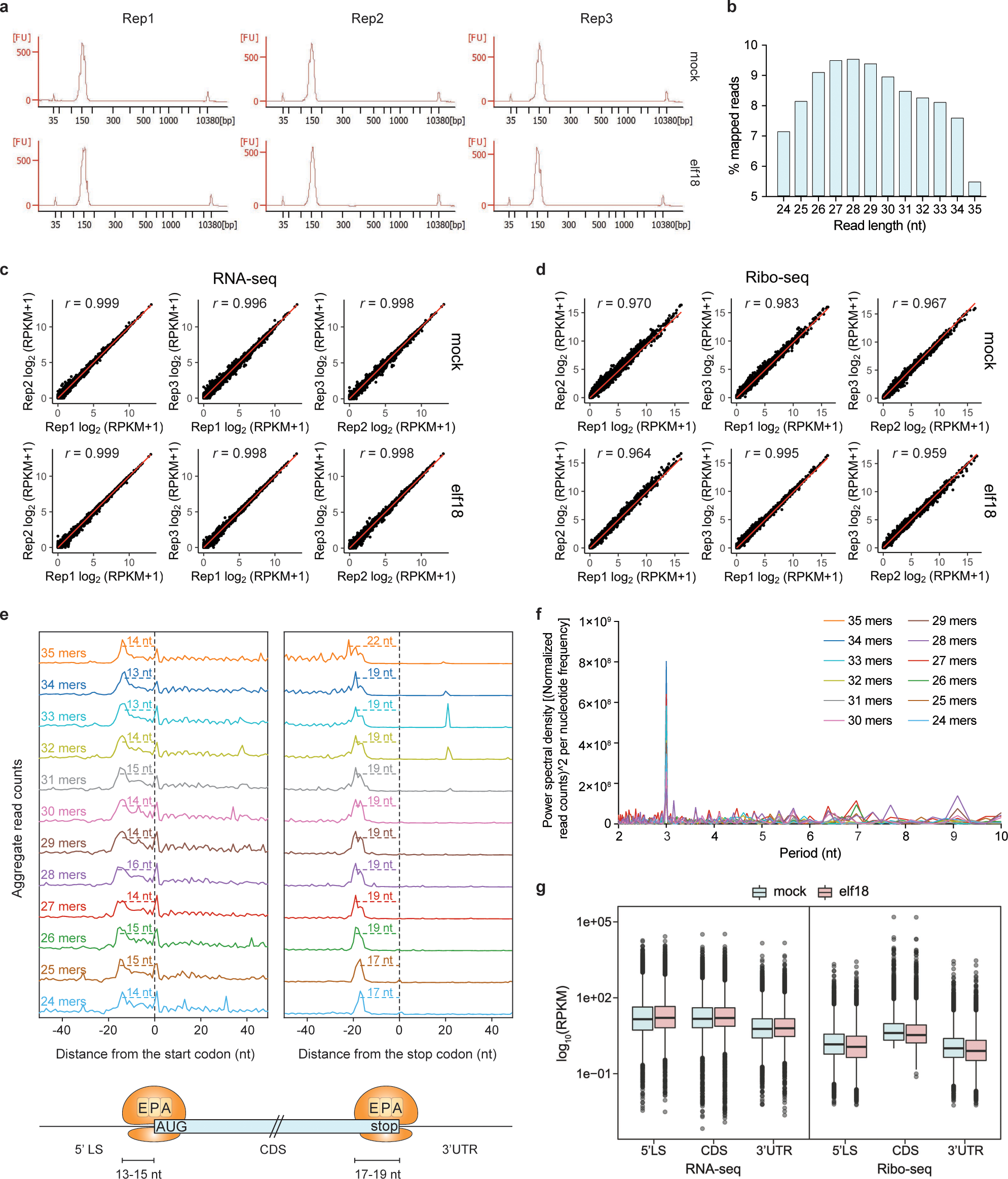
Quality and reproducibility of RNA-seq and Ribo-seq data. **a**, BioAnalyzer profiles showed high quality of the Ribo-seq libraries. Apart from the internal standard sized at 35 bp and 10380 bp, a single peak at ∼150 bp is present in all the libraries for mock and elf18 treatment in all three biological replicates (Reps 1-3). **b**, Length distribution of all reads from the Ribo-seq libraries. **c**, **d**, Correlations among the three replicates of RNA-seq (**c**) and Ribo-seq (**d**) data from mock- and elf18-treated samples. Data are shown as correlations of log_2_(RPKM+1) for all the genes. *r*, Pearson correlation coefficient. **e**, Metagene analysis on the average read counts surrounding start and stop codons for reads at different lengths, ranging from 35 nt (35 mers) (top) to 24 nt (24 mers) (bottom). P-site offsets were detected at the length of 13-15 nt surrounding start codons and at the length of 17-19 nt surrounding stop codons (bottom). 5’ LS, 5’ leader sequence. **f**, Power spectral density of normalized Ribo-seq read counts in the 300 nt window downstream of the start codon shows 3-nt periodicity. **g**, Total RNA-seq and Ribo-seq read distribution in 5’ LS, CDS, and 3’UTR. Boxes represent the interquartile range (IQR), and whiskers indicate data within 1.5 × IQR of the top (Q3) and bottom (Q1) quartiles. Grey circles represent RPKM values for individual outlier transcripts.

**Extended Data Fig. 2.**
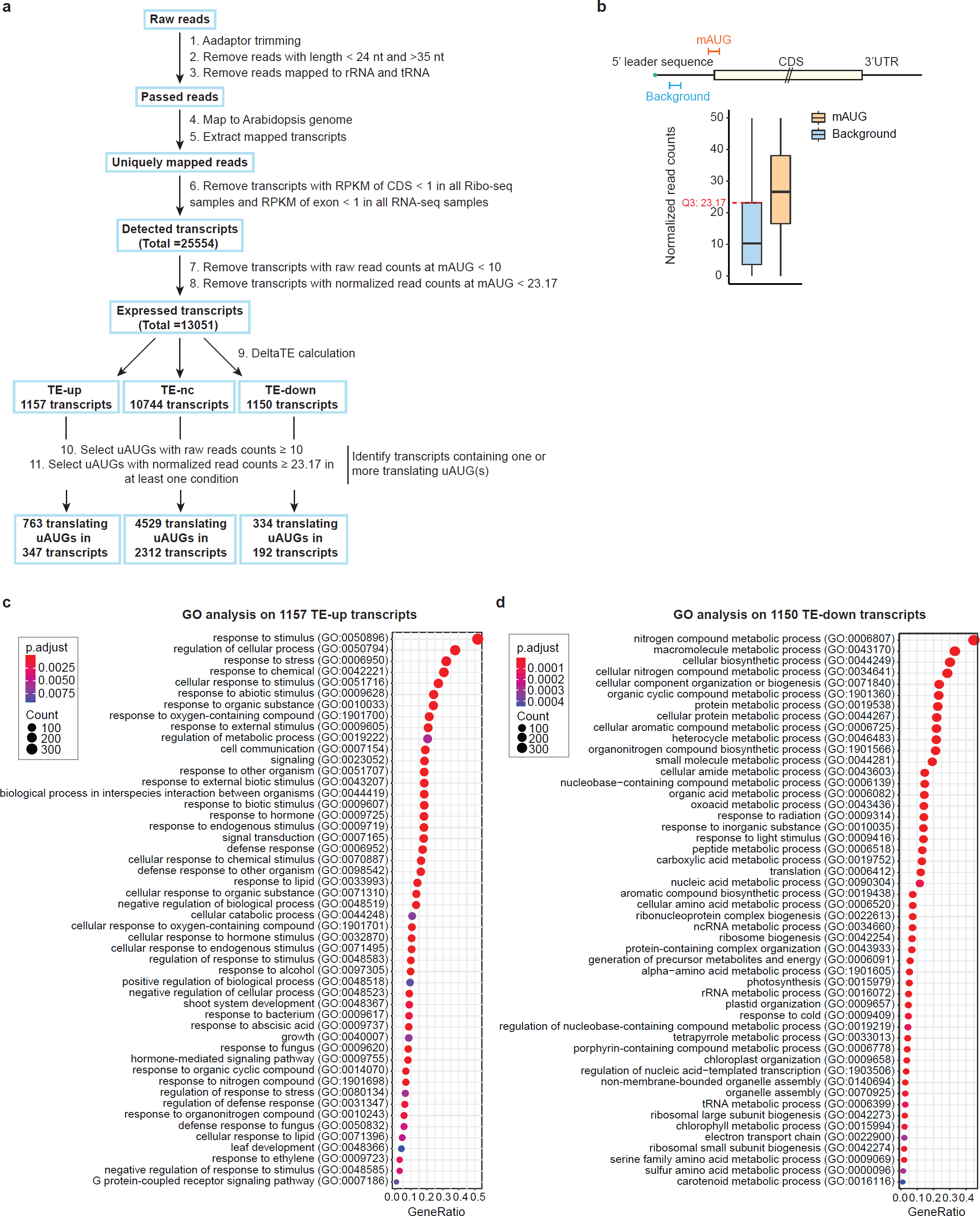
Global analysis of translational dynamics and uAUG-containing transcripts during pattern-triggered immunity. **a**, A flowchart of RNA-seq and Ribo-seq data analysis. **b,** Strategy for identification of translating mAUGs and uAUGs (see methods for details). **c, d,** Gene ontology (GO) analysis on the 1157 TE-up transcripts (**c**) and 1150 TE-down transcripts (**d**). The size of the dots represents the number of genes that fall into each group. The colour of the dots represents adjusted p-value.

**Extended Data Fig. 3.**
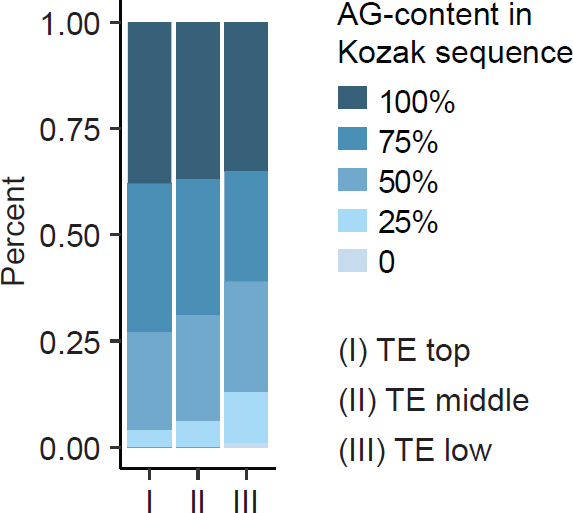
Comparison of Kozak sequence contexts surrounding mAUGs in trancripts with different levels of translational efficiency. Comparison of the AG-content in Kozak sequences flanking (I) top 100 transcripts with TE ranking the highest; (II) middle 100 transcripts with TE ranking 50%; (III) bottom 100 transcripts with TE ranking the lowest. Transcripts with similar abundance, 5’ leader sequence lengths and no uAUGs were selected for the comparison.

**Extended Data Fig. 4.**
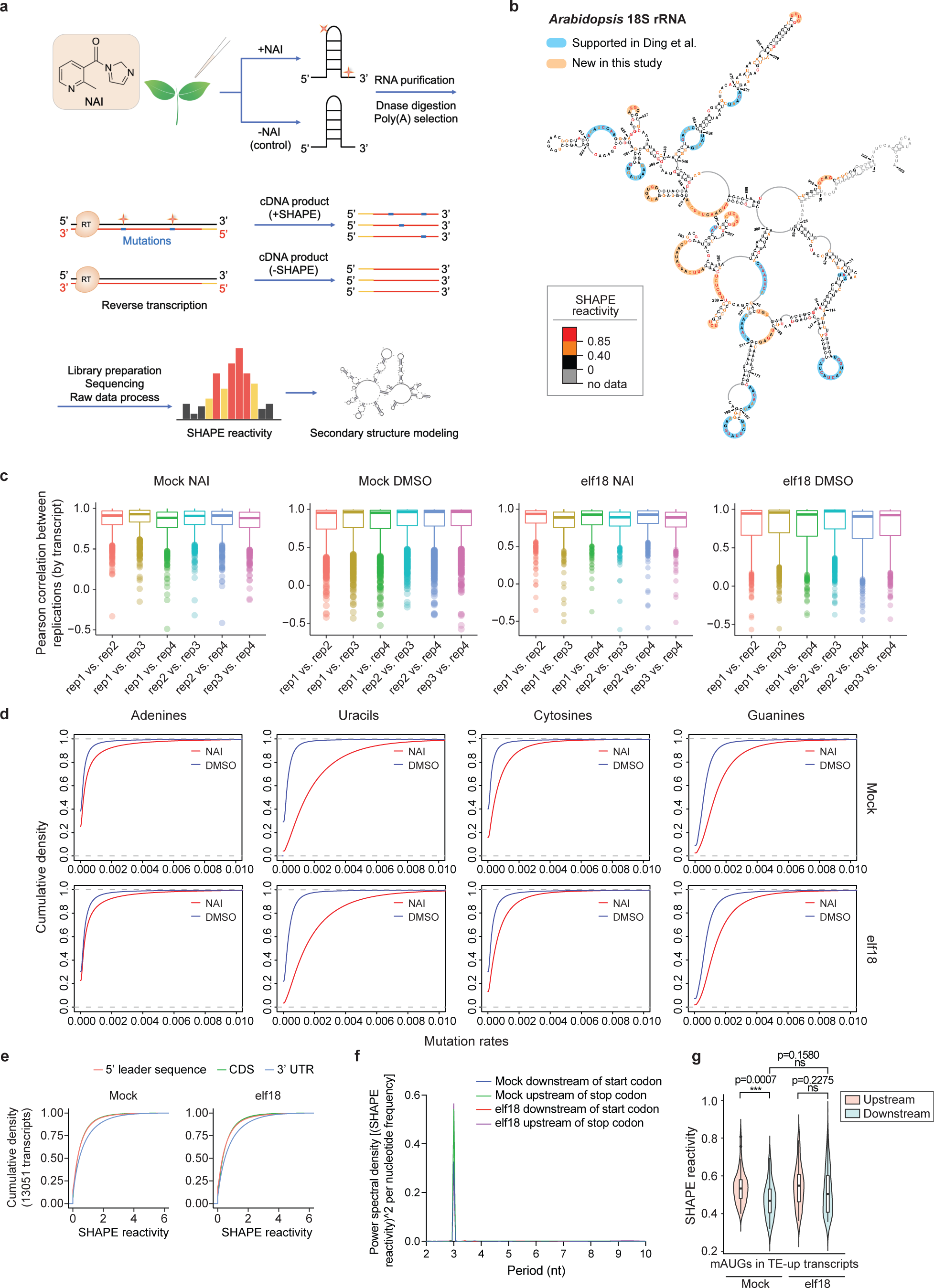
Quality and reproducibility of global and targeted *in planta* SHAPE-MaP. **a**, A flowchart of *in planta* SHAPE-MaP protocol. **b**, Comparison of *Arabidopsis* 18S rRNA secondary structure detected using the DMS-based structure-seq method performed by Ding et al.^39^ and the SHAPE-MaP protocol used in this study. **c**, Pearson correlation among the four SHAPE-MaP biological replicates (by transcript) under each treatment condition. Boxes represent the interquartile range (IQR), and whiskers indicate data within 1.5 × IQR of the top (Q3) and bottom (Q1) quartiles. Circles represent Pearson correlation values for outliers. **d**, Cumulative fraction on the mutation rates of every nucleotide under each treatment condition. **e**, Cumulative fraction on the SHAPE reactivity of 5’ leader sequence, CDS, and 3’UTR in mock and in response to elf18. **f**. Power spectral density of SHAPE reactivity shows 3-nt periodicity in the 150 nt window downstream of the start codon and in the 150 nt window upstream of the stop codon. **g**, Violin plot comparisons of SHAPE reactivities in the region 50 nt upstream or 50 nt downstream of mAUGs in the 347 TE-up transcripts in response to elf18 treatment. Boxes represent the interquartile range (IQR), and whiskers indicate data within 1.5 × IQR of the top (Q3) and bottom (Q1) quartiles. Circles represent values for outliers. Data were analysed by two tailed Mann-Whitney tests.

**Extended Data Fig. 5.**
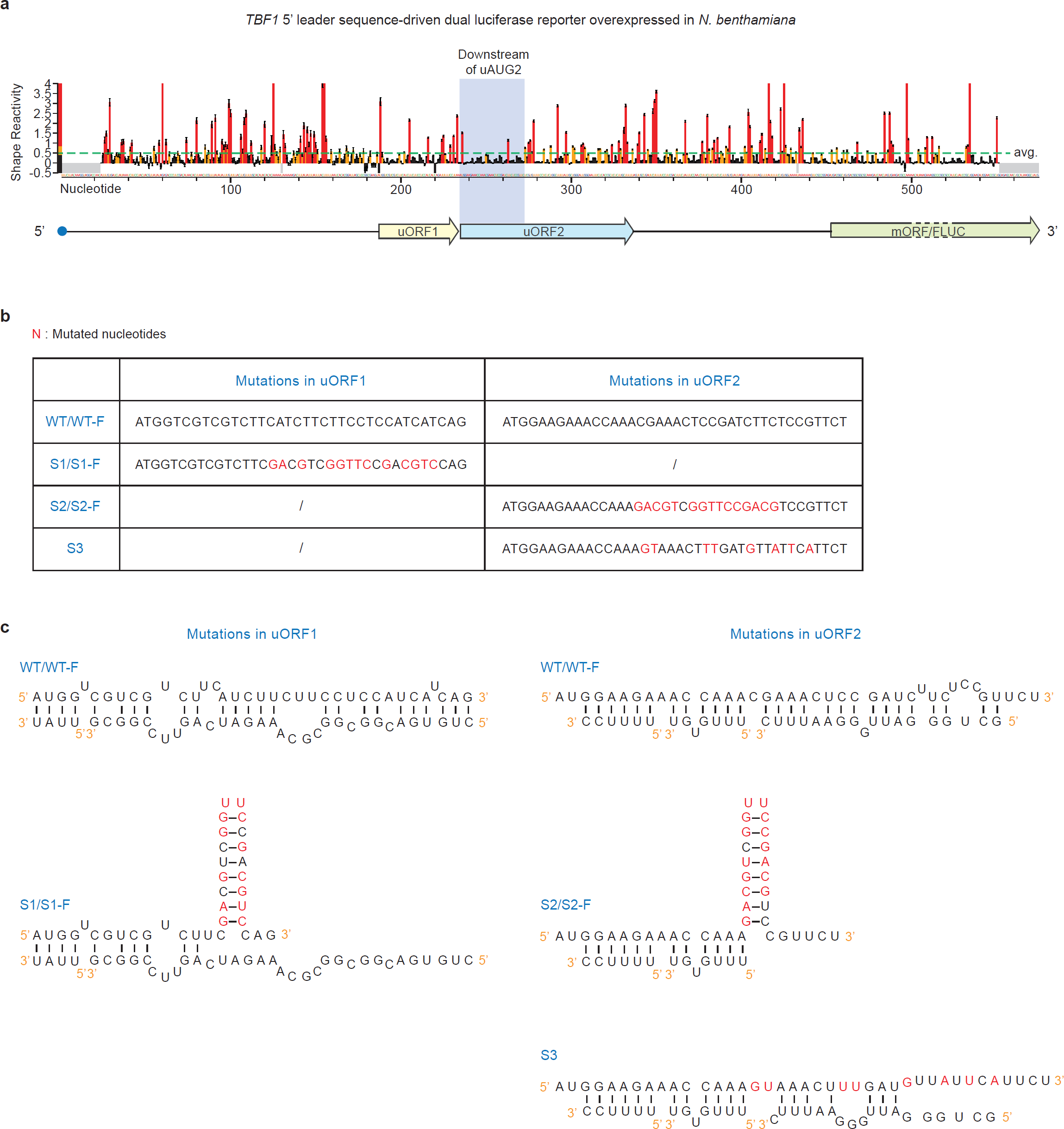
*In planta* SHAPE-MaP of the *TBF1* transcript. **a**, Overview of SHAPE reactivities across the 5’ leader sequence of the *35S:TBF1 5’ leader sequence-FLUC/35S:RLUC* dual-luciferase reporter overexpressed in *N.* benthamiana. The region 35 nt downstream of uAUG2 is highlighted in light blue. The green dashed line indicates average the SHAPE reactivity across the region. **b**, Mutagenesis on the secondary structures in the *TBF1* transcript. Red letters indicate mutated nucleotides. Notably, the mutated regions are not conserved in the primary protein sequence across Angiosperms^18, 27^. **c**, Effects of mutating the RNA secondary structures in the *TBF1* transcript. Structural models generated according to *in planta* SHAPE-MaP in the region downstream of uAUG1 and uAUG2 are shown on the left and right, respectively. In S1/S1-F, a stable hairpin (Δx1D43A; = -9.9) was introduced in the downstream region of uAUG1 to make this region more structured. In S2/S2-F, the same hairpin was introduced in the downstream region of uAUG2 to make this region more structured. In S3, some of the base pairings in WT was disrupted to make the region downstream of uAUG2 less structured.

**Extended Data Fig. 6.**
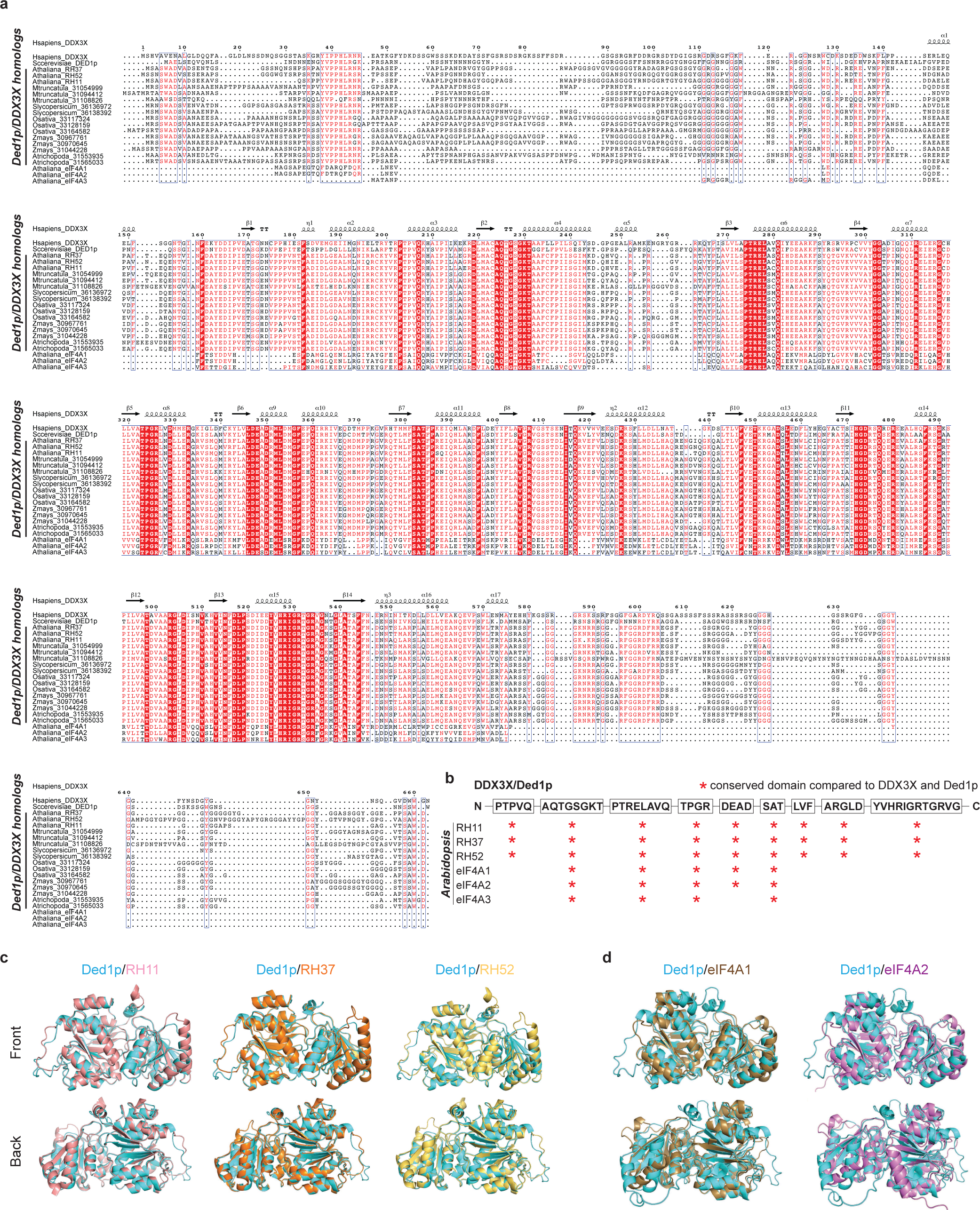
Structural similarities of *Arabidopsis* homologous RNA helicases RH11, RH37 and RH52 with yeast Ded1p and mammalian DDX3X. **a**, Protein sequence alignment of *Arabidopsis* RH11, RH37, and RH52 with their homologues in another five angiosperm species: *Amborella trichopoda* (Atrichopoda), *Zea mays* (Zmays), *Oryza sativa* (Osativa), *Solanum lycopersicum* (Slycopersicum), and *Medicago truncatula* (Mtruncatula), and together with yeast Ded1p, human DDX3X, as well as *Arabidopsis* eukaryotic translation Initiation Factor 4A (eIF4A) homologues. Numbers followed each name are PACIDs. ESPript 3.0^49^ was used for protein sequence alignment visualization. Human DDX3X secondary structure elements were used as reference. **b**, Domain conservation of *Arabidopsis* RH11, RH37, RH52, eIF4A1, eIF4A2, and eIF4A1 with DDX3X/Ded1p regarding the nine sequence motifs (in the boxes and are illustrated from N terminus to C terminus). Conserved domains are indicated with red asterisks. **c**, **d**, Pairwise alignment of yeast Ded1p with *Arabidopsis* RH11, RH37, and RH52 (**c**) and with *Arabidopsis* eIF4A1 and eIF4A2 (**d**) shows that RH11, RH37 and RH52, but not eIF4A1 and eIF4A2, are structurally similar to Ded1p. Protein structures were predicted by AlphaFold^44^.

